# The unique tropism of *Mycobacterium leprae* to the nasal epithelial cells can be explained by the mammalian cell entry protein 1A

**DOI:** 10.1101/373324

**Authors:** Viesta Beby Fadlitha, Fuki Yamamoto, Irfan Idris, Haslindah Dahlan, Naoya Sato, Vienza Beby Aftitah, Andini Febriyanda, Takao Fujimura, Hiroaki Takimoto

## Abstract

Leprosy is a chronic infection where the skin and peripheral nervous system is invaded by *Mycobacterium leprae*. The infection mechanism remains unknown in part because culture methods have not been established yet for *M.leprae.*

Mce1A protein (442 aa) is coded by mce1A (1326 bp) of *M.leprae*. The mce1A homolog in *Mycobacterium tuberculosis* is known to be associated with *M.tuberculosis* epithelial cell entry, and survival and multiplication within macrophages. Studies using recombinant proteins have indicated that mce1A of *M.leprae* is also associated with epithelial cell entry. This study is aimed at identifying particular sequences within mce1A associated with *M.leprae* epithelial cell entry.

Recombinant proteins having N-terminus and C-terminus truncations of the mce1A region of *M.leprae* were created in *Eschericia coli.* Entry activity of latex beads, coated with these truncated proteins (r-lep37kDa and r-lep27kDa), into HeLa cells was observed by electron microscopy. The entry activity was preserved even when 315 bp (105 aa) and 922 bp (308 aa) was truncated from the N-terminus and C-terminus, respectively. This 316 – 921 bp region was divided into three sub-regions: 316 – 531 bp (InvX), 532 – 753 bp (InvY), and 754 – 921 bp (InvZ). Each sub-region was cloned into an AIDA vector and expressed on the surface of *E.coli.* Entry of these *E.coli* into monolayer-cultured HeLa and RPMI2650 cells was observed by electron microscopy. Only *E.coli* harboring the InvX sub-region exhibited cell entry. InvX was further divided into 4 domains, InvXa - InvXd, containing sequences 1 – 24 aa, 25 – 46 aa, 47 – 57 aa, and 58 – 72 aa, respectively.

Recombinant *E.coli*, expressing each of InvXa - InvXd on the surface, were treated with antibodies against these domains, then added to monolayer cultured RPMI cells. The effectiveness of these antibodies in preventing cell entry was studied by colony counting. Entry activity was suppressed by antibodies against InvXa, InvXb, and InvXd. This suggests that these three InvX domains of mce1A are important for *M.leprae* invasion into nasal epithelial cells.

**Author Summary:** Mce1A protein is encoded by the mce1A region of mce1 locus of *M.tuberculosis* and *M.leprae*, and is involved in the bacteria’s invasion into epithelial cells. The present study revealed that the active sequence of *M.leprae* involved in the invasion into nasal mucosa epithelial cells is present in the 316-531 bp region of mce1A.

The most important region of mce1A protein involved in the invasion of *M.tuberculosis* into human epithelial cells is called the InvIII region, which is located between amino acids at position 130 to 152. The InvIII region of *M.tuberculosis* corresponds to InvXb of *M.leprae*. The sequences of these regions are identical between amino acids at positions 10 to position 22 as counted from the N terminus, except that amino acids at positions 1 to 3, 5, 8, 9, 13 are different between *M.leprae* and *M.tuberculosis*. Suppression test results also indicated that the most important region of mce1A protein of *M.leprae* involved in the invasion into human epithelial cells is different from that *M.tuberculosis*. While *M.tuberculosis* has 3,959 protein-encoding genes and only 6 pseudogenes, *M.leprae* has only 1,604 protein-encoding genes but has 1,116 pseudogenes indicating that in *M.leprae*, far more proteins are inactivated as compared to *M.tuberculosis*. The present study also revealed that, as in *M.tuberculosis,* the mce1A protein is expressed on the surface of bacteria as a native protein. In light of these data, the mce1A protein is considered to be one of the most important proteins involved in the invasion of *M.leprae* into nasal mucosa epithelial cells.

## Introduction

Hansen’s disease is a chronic infection with acid fast bacillus where skin and peripheral nerves are damaged by the infection with *Mycobacterium leprae* (*M.leprae*). Although the number of Hansen’s disease cases has drastically decreased in developed countries, worldwide, the number of new cases of Hansen’s disease has only dipped below 200,000 per year. Hansen’s disease is one of the Neglected Tropical Disease (NTDs) and is still a major problem against public health[1].

Hansen’s disease can be broadly divided into tuberculoid leprosy (T type) and lepromatous leprosy (L type), depending on the host immune response to *M.leprae*[2]. Tuberculoid leprosy triggers predominantly cellular immunity response, and is also called paucibacillary, because very few are detected at the focus of infection or nasal mucosal membrane. On the other hand, lepromatous leprosy triggers predominantly humoral immunity, and is also called multibacillary, because it is detected in a large amount at the focus of infection and, in particular, from nasal mucosal membrane. Nasal discharge from lepromatous leprosy patients, therefore, is considered as the main source of the infection[3]. Infection of Hansen’s disease has conventionally been considered to occur through close skin contact or through wounds, but recently another infection mode, in which *M.leprae* in the aerosol from nasal discharge of lepromatous leprosy patients invades into the upper respiratory tract and nasal mucosal membrane to cause infection, has come to be recognized[3-10]. However, the invasion mechanism in this infection mode has not been extensively studied yet.

*M.leprae* cannot be artificially cultured. One possible reason for this is the presence of a large number of pseudogenes. *M.leprae* has various enzyme-coding genes that are replaced with pseudogenes, and therefore has only a minimum metabolic activity and multiplies in macrophages and Schwann cells. Invasion mechanism of *M.leprae* into Schwan cells have been studied by Rambukkana, et al., in details. The study revealed that laminin-2 present in the basal lamina surrounding Schwann cells serves as a receptor, and histone-like protein Hlp/LBP expressed on the surface of the bacteria binds to phenolic glycolipid PGL-1, making the entry into the cell possible[11-14].

To infect Schwann cells, *M.leprae* has to invade the epidermal cells first. The mechanism of *M.leprae* invasion into the epidermal cells, however, has not been elucidated yet. Meanwhile, gene regions involved in the invasion of *Mycobacterium tuberculosis (M.tuberculosis)* into epidermal cells are already known[15,16]. Casali et al. reported that, using adhesin involved in diffuse adherence (AIDA) method, the region coded for by 316 - 531 bp of *M.tuberculosis* mce1A region (Rv 0169; 198534 - 199898 bp, 1365 bp) is expressed on the surface of *E.coli* as a polypeptide chain, thereby imparting the *E.coli* with the ability to invade HeLa cells, that the invasion activity is inhibited by the monoclonal antibody that recognizes the continuous peptide of InvIII region (388 – 453 bp)[17,18]. It became clear that *M.leprae* includes a region (ML2589, 1326 bp) highly homologous to mce1A protein of *M.tuberculosis.* Sato et al. reported that a recombinant protein, a 37 kDa protein encoded by 73 - 921 bp, which is the mce1A region excluding the signal sequence, was found to have an invasion activity into epidermal cells[19]. However, the active sequence involved in the invasion by *M.leprae* into epidermal cells has not been identified. The present study was conducted to identify the active sequence in the mce1A region. In this study, the N-terminus and C-terminus truncated proteins expressed on the *E.coli,* where *E.coli* with specific regions are expressed thereon by the AIDA method, and hyperimmune antisera against the invasion region are used to investigate the invasion activity into epidermal cells.

## Material and methods

### Bacterial strains and plasmid

The genomic DNA used in the study was isolated from *M.leprae* strain Thai 53, which was maintained at Leprosy Research Centre, National Institute of Infectious Diseases, Japan, as previously described[20,21]. The pQE30 plasmid and *E.coli* M15 (pREP4) were purchased from Qiagen (Valencia, CA). The pQE30 plasmid was used as expression vector. *E.coli* M15 (pREP4) was used as a host for the vector, as recommended by the manufacturer.

The pMK90 plasmid and *E.coli* UT4400 were obtained from Dr. Riley (University of California at Berkeley, California, USA).

### Construction of vector

In Sanger Center *M. leprae* strain TN complete genome sequence, mce1A gene is a 1326 bp putative ORF located between positions 3092446 and 3093771 (NCBI-GeneID: 910890). The mce1A DNA sequence of strain Thai 53 was identical to that of strain TN. It was subcloned into pQE30 vector in a truncated reading frame. The 603 bp ORF deleted at 5’ and 3’ ends of mce1A gene is located between positions 316 and 921 (Fig 1). This sequence was amplified by polymerase chain reaction (PCR) directly from the genomic DNA of *M. leprae* strain Thai 53 with oligonucleotide primers designed to introduce SacI and HindIII endonuclease restriction sites at the ends. The amplified products were ligated into the pQE30 vector linearized with SacI and HindIII. The use of pQE30 vector allowed the plasmid to express the mce1A product with a polyhistidine (6 x His) tag at the N-terminus (r-mce1Ap). The resultant plasmid was cloned into *E.coli* M15 (pREP4) by electroporation (Gene Pulser II, Bio-Rad, Hercules, CA), according to the manufacturer’s instructions.

**Fig 1.**
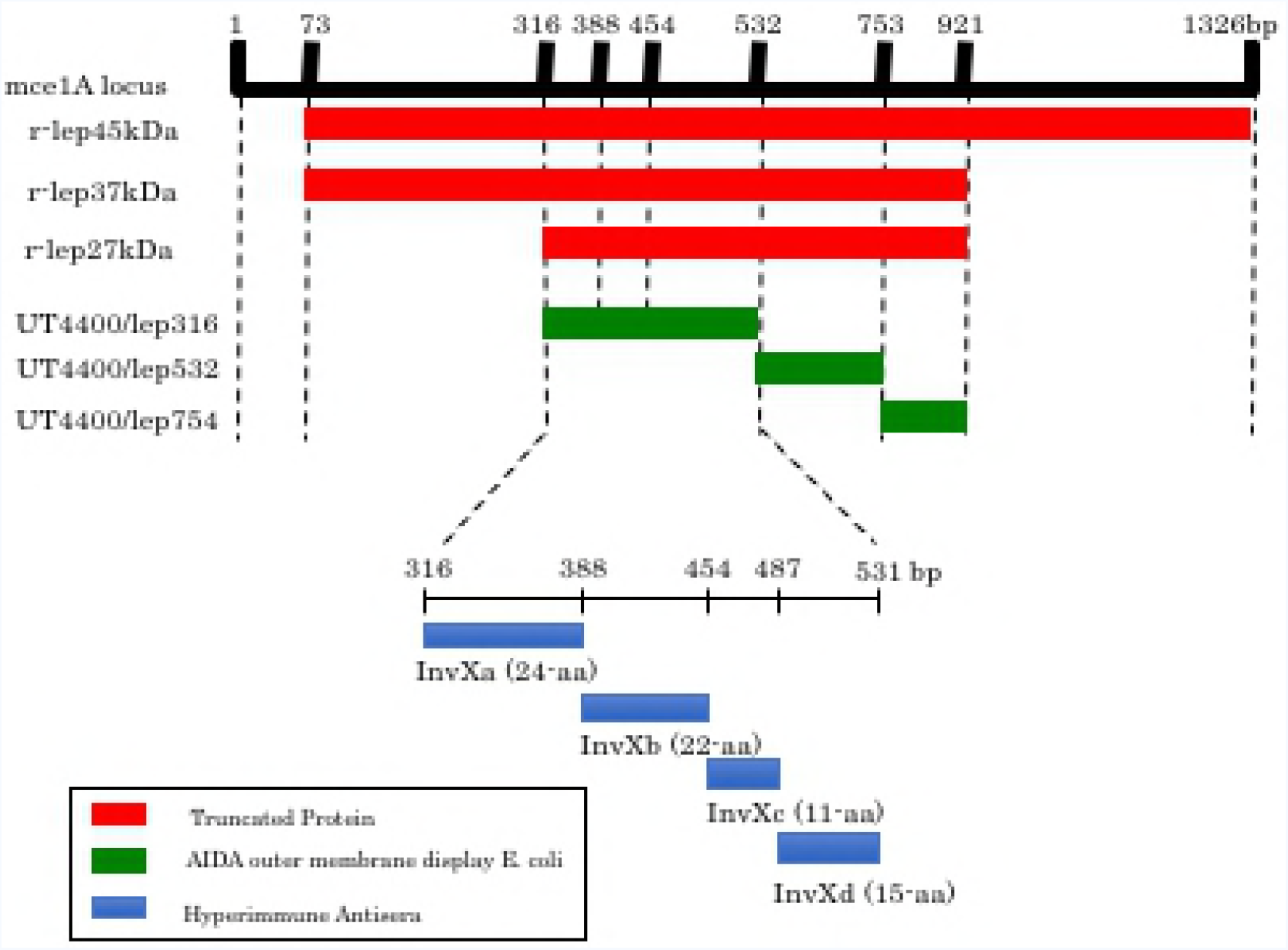
r-mce1A truncated protein regions expressed on *E.coli*, regions externally expressed using AIDA method, and regions for which hyperimmune antisera is prepared. Using r-lep45kDa protein encoded in the entire region of mce1A minus signal sequences (73 – 1326 bp) as a reference, the protein with its C terminus truncated to 921 bp was labeled as r-lep37kDa protein, and the protein with its N terminus truncated to 315 bp was labeled as r-lep27kDa protein. Furthermore, 316 – 921 bp region was divided into 316 – 531 bp, 532 – 753 bp, and 754 – 921 bp, and recombinant *E.coli* in which these regions are externally expressed using AIDA method were prepared. InvX region (316 – 531 bp) was further divided into InvXa (316 – 387 bp), InvXb (388 – 453 bp), InvXc (454 – 486 bp), InvXd (487 – 531 bp), and hyperimmune antisera for each region was prepared.

Plasmid pMK90 is an ampicillin-resistant pBR322 derivative that expresses a recombinant AIDA protein under the control of its own promotor[22]. The AIDA coding sequence has been altered to remove the native passenger; it consists of a 49-amino-acid signal peptide. A 78-amino-acid linker region incorporating a multiple cloning site, and the entire 440-amino-acid barrel core. A 216 bp DNA fragment encoding InvX (*M.leprae* positions 3092761 – 3092976 bp), 222 bp DNA fragment encoding InvY (*M.leprae* positions 3092977 – 3093198 bp), 168 bp DNA fragment encoding InvZ (*M.leprae* positions 3093199 – 3093366 bp) was amplified by PCR from a plasmid containing mce1A and cloned into pMK90, generating pMK100. The correct insert was confirmed by sequencing. The amino acid sequence of InvX is VNADIKATTVGGKYVSLTTPPEHPSQKRLTPQTVIDARSVTTEINTLFQTITLIAEKVDPIKLNLTLSAAAQ (316 – 531 bp), the amino acid sequence of InvY is SLAGLGERFGQSIVNGNSVLDDVNSQLPQARHDIQQLASLGDTYANSASDFFDFLNN SIVTSRTI (532 – 753 bp), and the amino acid sequence of InvZ is VLLAAVGFGNTGADIFSRSGPYLARGAADLVPTAQLLDTYSPAIFCTLRNYHDIEP (754 – 921 bp) (Fig 1).

### Protein expression and purification

Recombinant protein was expressed and purified according to manufacture’s instruction. Briefly, *E.coli* M15 [pREP4] containing pQE30/mce1A plasmid was grown overnight in 10-ml superbroth containing 100 μg/ml ampicillin and 50 μg/ml kanamycin. A 500 μl aliquot of bacterial suspension was pelleted, resuspended in 30 ml of superbroth and incubated at 37°C for 1 – 2 h until OD_600_ = 0.6. Then isopropyl β-D-thiogalactoside was added to final concentration 1 mM and incubated for 3 h at 37°C. The induced and uninduced r-*E.coli* strains were analyzed by SDS-polyacryamide gel electrophoresis (SDS-PAGE). The newly expressed protein formed an inclusion body in the r-*E.coli* host. The inclusion body was therefore purified under denaturing conditions according to the instructions of the expression vector’s respective manufactures. The 6 x His tag mce1A solubilized with lysis buffer (6 M Guanidine, 10 mM Tris—HCl, 100 mM NaH_2_PO_4_, pH 8.0) was bound to a Ni—NTA resin column equilibrated with lysis buffer, and was eluted by elusion buffer (6 M Guanidine, 10 mM Tris—HCl, 100 mM NaH_2_PO_4_, 20—250 mM imidazole, pH 6.3). The eluted protein were subsequently refolded with 1 mM dithiothreitol (Sigma, St. Louis, MO, USA) and 0.1 mM phenylmethylsulfonyl fluoride (Sigma) by dialysis, gradually removing guanidine. The r-mce1A was finally purified and refolded as a soluble protein. The protein was separated by SDS-PAGE and was analyzed for purity by Coomasie brilliant blue R-250 staining.

### Immunoelectron microscopy

The antibody against the mce1A protein was prepared in Balb/c mice (a 45 kDa recombinant mce1A protein prepared previously using *E.coli* was used as the immunogen). The r-45kDa-mce1A protein was used for antibody production because it was most abundantly expressed in the *E.coli* host that we used. The r-45kDa-mce1A protein was mixed with Titer Max Gold (AdipoGen Life Sciences, Liestal, Switzerland) of the same amount. Approximately 100 μg of the protein was administered subcutaneously at five sites in four 7-weeks-old Balb/c mice, followed by two booster injections of 100 μg each 2 and 4 weeks after the first injection. Blood was harvested 2 weeks after the last booster dose.

The antiserum was used in the colloidal gold immunoelectron microscopy. A bacterial pellet (containing ≈ 10^7^ organisms) of *M.leprae* strain Thai 53 that that had been multiplied in footpads of athymic nude mice, was fixed in 3% glutaraldehyde in phosphate buffer saline (PBS) pH 7.6 for 24 h, washed five times in PBS and then exposed at 4°C for 16 h to a 1:1000 dilution of the mice antibody raised against mce1A. The suspension was then washed and incubated at 4°C for 16 h with colloidal gold suspension containing 5 nm gold particles (1.9 × 10^13^ particles ml^−1^) conjugated to anti-mice IgG goat antibody (Amersham/GE Health Care Life Science, Tokyo, Japan). The cells were washed again five times in PBS, stained with 0.1% uranyl acetate in water and examined with a HITACHI model H-15 electron microscope.

### Cell Culture

HeLa cells and RPMI2650 cells were purchased from America Type Culture Collection (ATCC, Manassas, VA). HeLa cells (ATCC CCL-2) were maintained with Dulbecco’s modified Eagle’s media (DMEM; Invitrogen, Carlsbad, CA) supplemented with 50 μg/ml gentamicin (GM) and 10% fetal bovine serum (FBS) (JRH Bioscience, Lenexa, KS). RPMI2650 human epithelial nasal septal cell line (ATCC CCL30) was grown in Eagle’s minimum essential medium (EMEM; Invitrogen, Carlsbad, CA) supplemented with 50 μg/ml GM and 10% FBS. Cells were maintained in culture and for the assay, were detached from the plastic by using 0.25% Trypsin-EDTA (1x) with phenol red (Gibco, Grand Island, NY, USA) at 37°C. The cells were then centrifuged at 280 × g for 7 min at 4°C, counted in Neubauer hemocytometer, and plated into tissue culture well or flask at 37°C in a 5% CO_2_ atmosphere.

### Cell uptake assay of protein-coated latex beads by electron microscopy

A 30 μ of stock suspensions of 1.1 μm diameter polystyrene latex beads, containing 5 × 10^8^ beads/ml (Sigma), were mixed in 150 μl of PBS containing 50 μg/ml of each set of protein and incubated for 16 h at 37°C. After incubation, the samples were centrifuged at 7000 × g and resuspended in 750 μl of PBS. A 500-μl sample of this suspension was added to a near-confluent cultured cell monolayer grown in a 25-cm^2^ flask containing 7 ml of appropriate media for cultured cells. The cells were incubated for 5 h at 37°C in a CO_2_ incubator, washed four times with PBS and one time with 0.1 M cacodylate phosphate buffer (pH 7.6), and then collected with cell-scraper (Becton Dickinson, Japan). The collected cells were fixed with 2% glutaraldehyde in 0.1 M cacodylate phosphate buffer (pH 7.6) at 4°C overnight, post-fixed with 1% osmium tetroxide in PBS, dehydrated through graded ethanol solutions and embedded in Spurr’s low-viscosity embedding media. The ultrathin sections were stained with uranyl acetate and lead citrate and examined with a JEM-1200EX (JEOL, Tokyo, Japan) transmission electron microscope. Coated beads with bovine serum albumin (BSA) fraction V (Boehringer Mannheim, GmbH, Germany) were used as negative controls.

### Immunofluorescence microscopy

*E.coli* cells were fixed onto microscope slides with 0.4% paraformaldehyde for 10 min at room temperature, and non-specific binding was blocked by incubation in 1% (wt/vol) BSA for 30 min. Slides were incubated for 1 h with a 1:200 dilution of a rabbit antibody raised against InvXa, InvXb, InvXc, and InvXd washed, and incubated with 1:1000 dilution of fluorescein isothiocyanate-labeled anti-rabbit antibody (Abcam Plc, Cambridge, UK) for 30 min. After extensive washing, the coverslips were mounted. Slides were viewed on an Olympus BX51 inverted microscope with an epifluorescence attachment.

### Invasion assay

#### InvX, InvY, InvZ mediated invasion of *E.coli* into the nasal epithelial cells by electron microscopy

The culture medium of a monolayer culture of 1.15 × 10^6^ HeLa cells and 3 × 10^7^ RPMI2650 cells, was replaced with a culture medium that does not contain antibiotic substances. Then *E.coli* externally expressing by AIDA (UT4400/lep316, UT4400/lep532, UT4400/lep754) was added at a bacteria to cells ratio of 100:1, and incubated in a CO_2_ incubator at 37°C, for 9 h for HeLa cells and 6 h for RPMI2650 cells. After culturing, the cell surface was washed with PBS, and then harvested using a cell scraper.

Infected cells were prepared for examination by transmission electron microscopy as previously described^[16]^. Briefly, cells were fixed in 2% glutaraldehyde and stained with osmium tetroxide solution before dehydration through graded ethanol solutions. Cells were embedded in Spurr’s low-viscosity embedding medium, ultrathin sections were stained with uranyl acetate and lead citrate. Samples were examined with a JEM-1230 (JEOL) transmission electron microscope.

### Gentamicin protection assays

Gentamicin protection assays were performed according to the method of Elsinghorst[23]. RPMI2650 cells were seeded at 5 × 10^5^ cells per well directly into 24-well plates and cultured for 24 h until confluent. Cell culture medium was modified to contain no antibiotics. Recombinant *E.coli* cells were added to the monolayer at a multiplicity of infection (MOI) of 10:1 and incubated at 37°C for 3 h. To enumerate intracellular bacteria, the monolayer was washed five times with PBS and incubated with medium containing 100μg of GM (Sigma) per ml for 2 h to kill extracellular bacteria and permit the enumeration of intracellular bacteria. The monolayer was again washed five times with PBS and lysed with 0.1% Triton X-100 (Eastman Kodak, Rochester, NY). Serial dilutions of released bacteria were plated for counting. Results shown are the mean values for an experiment performed in triplicate. Each experiment was performed three times using independent cultures, with similar result.

### Inhibition Assay

InvXa, InvXb, InvXc, and InvXd antibodies were added in the amount of 1/200 (200 μg) to *E.coli* that externally express UT4400/lep316 by AIDA adjusted to 1 × 108 CFU/ml. This was allowed to react on a rotating platform at 4°C overnight to make antibody-treated bacteria. *E.coli* externally expressing proteins by AIDA were allowed to react with IgG from healthy, control rabbits in a similar manner as the control, where this was used as the bacteria untreated by antibody. After the medium for RPMI2650 cells, which were monolayer-cultured in a 24- well plates, 5 × 10^5^ cells/well, was replaced with a medium not containing antibiotic agent, the antibody-treated bacteria and untreated bacteria were added at a bacteria to cells ratio of 30:1. After culturing in CO_2_ incubator at 37°C for 3 h, the surface of the cells were washed with PBS five times, and the medium was replaced with a 100 μg/ml GM-appended DME medium to kill the bacteria outside the cells, followed by additional incubation for 2 h. The surface of cells was washed with PBS, and then 0.1% Triton X-100-added PBS was added in the amount of 1 ml/well to break the cells and the bacteria inside the cells were harvested. The harvested bacteria suspension liquid was serially diluted 10 times with PBS, and then was applied to Heart Infusion agar medium (Nissui, Tokyo, Japan). This was left overnight at 37°C, and then the colonies were counted to determine the number of bacteria entered into the cells. The cultured cells were prepared in the amount of 3 wells each, and the average of each well and standard deviation were calculated and the result was presented on a graph.

### Ethics statement

This study was approved by the Institutional Animal Care and Use Committee (Permission number: 2013153) and carried out in accordance with the KITASATO University Animal Experimentation Regulations.

## Result

### Immunoelectron microscopy of *M.leprae*

Immunoelectron microscopy was employed to determine whether recombinant *E.coli* expressed the mce1A protein on the cell surface.

The bacilli expressing mce1A protein were pretreated with an antibody (Ab) raised against r-45 kDa mce1A protein, and were followed by incubation with anti-IgG Ab-conjugated colloidal gold particles (Fig 2). The immunoelectron microscopic study revealed that the native mce1A protein is expressed on the surface of bacilli. This confirms that the recombinant *E.coli* not only expresses mce1, but the mce1 is transported to the cell surface and sufficiently presented such that it can bind the antibody against it.

**Fig 2.**
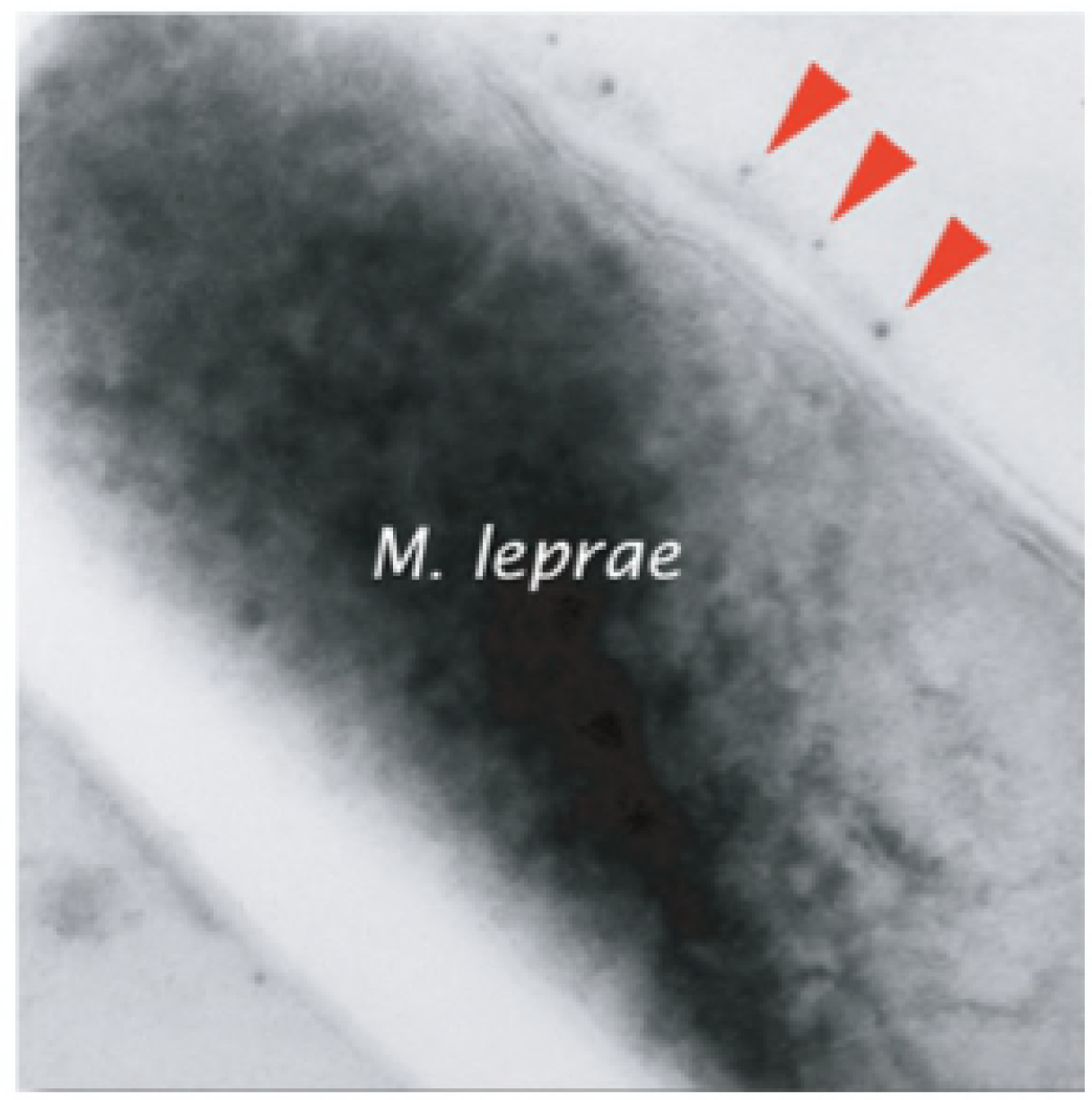
Colloidal gold immunoelectron microscopic analysis of *M.leprae* strain Thai 53. The bacilli were pretreated with an antibody (Ab) raised against r-45 kDa mce1A protein, and were followed by incubation with anti-IgG Ab-conjugated colloidal gold particles. Gold particles are shown decorating the surface of the *M.leprae* bacillus (arrowheads).

### HeLa cell uptake assay of protein-coated polystyrene latex microbeads by electron imcroscopy

The active sequence involved in the invasion into the epithelial cells was investigated in the following manner. The r-lep37kDa protein, which had been prepared in the previous experiment using r-lep45kDa protein as the reference by truncating the C terminus to 308 aa (922 bp), was further truncated to 105 aa (315 bp) from N terminus to provide r-lep27kDa protein where the proteins using were expressed using an *E.coli* expression system (Fig 1). Each of the truncated protein was observed for invasion activity into HeLa cells using an electron microscope. In this observation, images of beads coated with r-lep37kDa protein and beads coated with r-lep27kDa protein invading into the cytoplasm of HeLa cells were captured, but BSA-coated beads, which are the negative control, were not found to invade the cytoplasm (Fig 3). This result suggest that the active sequence is present between 316 – 921 bp, which encodes r-lep27kDa protein.

**Fig 3.**
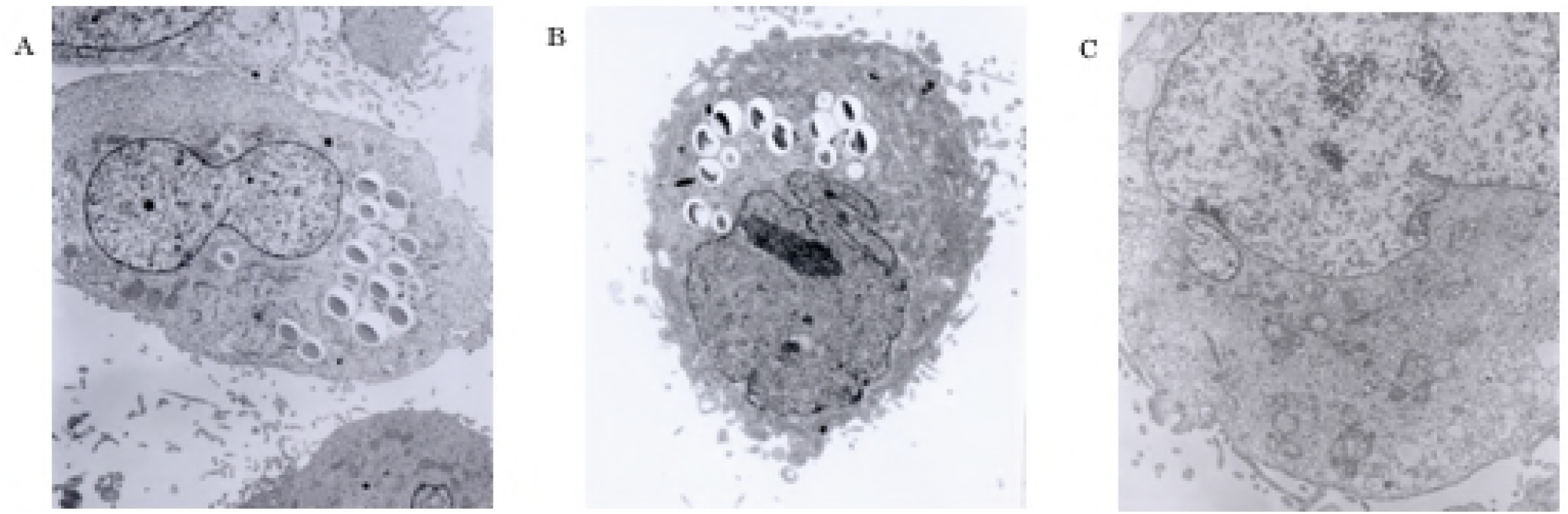
HeLa cells uptake assay of protein-coated polystyrene latex microbeads by electron microscopy. Monolayer-cultured HeLa cells and the truncated protein-coated beads and BSA-coated beads were allowed to react for 5 h, and the entry of beads into HeLa cells were observed under an electron microscope. As shown in the arrow, (A) r-lep37kDa protein-coated beads and (B) r-lep27kDa protein-coated beads were observed to enter into HeLa cells, but (C) BSA-coated beads were not observed to enter into HeLa cells. Magnification ×3500 (A), ×5000 (B,C)

### InvX, InvY, InvZ mediated invasion of *E.coli* into the nasal epithelial cells by electron microscopy

The active sequence was further inverstigated. The 316 – 921 bp region was divided into InvX; 316 – 531 bp, InvY: 532 – 753 bp, InvZ: 754 – 921 bp, and each of the regions was incorporated into AIDA vector to produce a recombinant *E.coli* externally expressing the proteins (Fig 1). The *E.coli* externally expressing the proteins by the AIDA method were observed for invasion activity into HeLa cells and RPMI2650 cells under the electron microscope. *E.coli* expressing InvX (UT4400/lep316) was found in abundance in the cytoplasm. *E.coli* expressing InvY and InvZ (UT4400/lep532 and UT4400/lep754) and UT4400 were observed present around the cells but not inside the cytoplasm (Fig 4). These results suggest that the active sequence is present in 316 – 531 bp (InvX).

**Fig 4.**
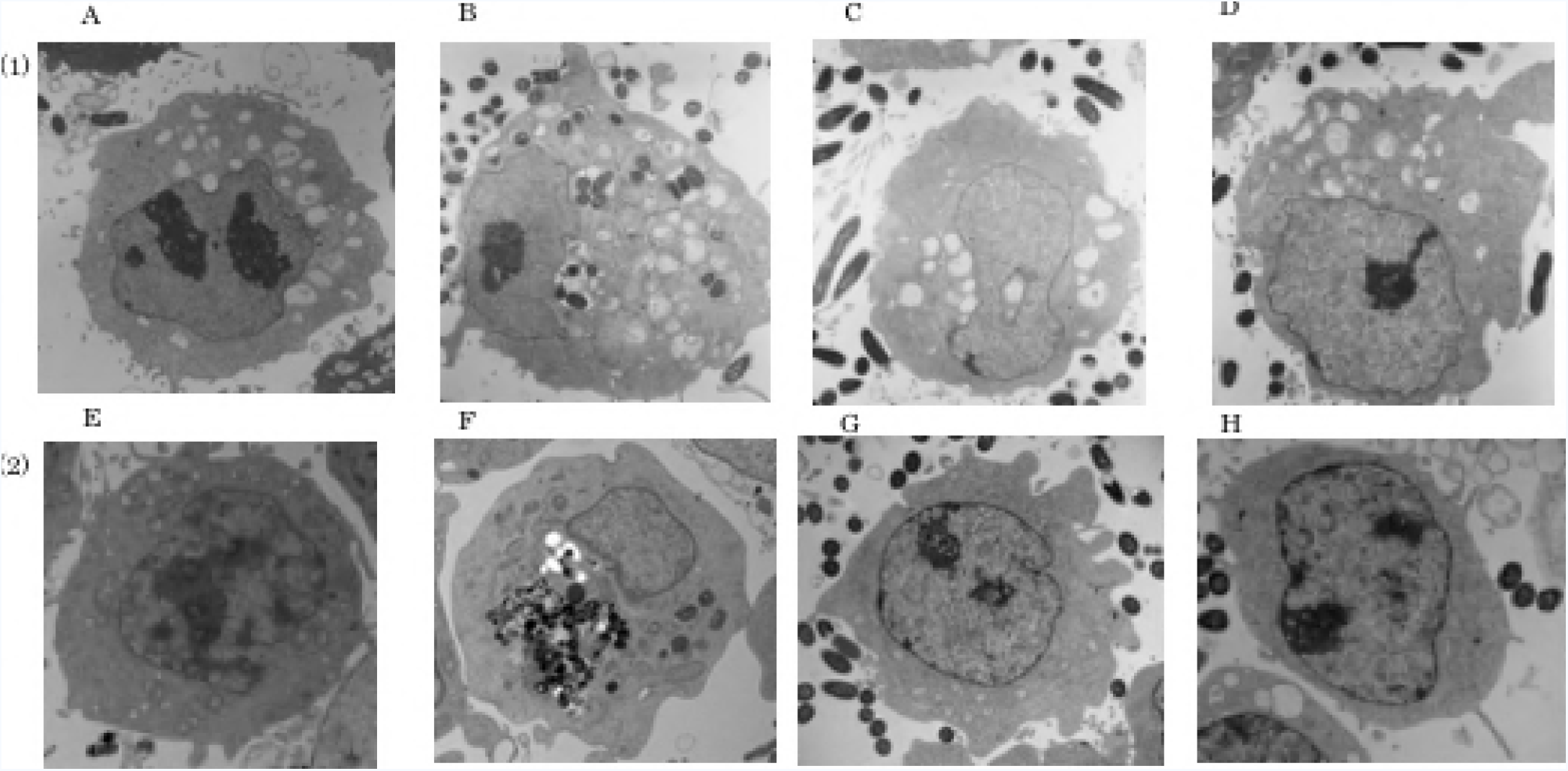
Transmission electron microscopy of InvX, InvY, InvZ mediated invasion of *E.coli* into HeLa and nasal epithelial cells. UT4400 (A,E), UT4400/lep316 (B,F), UT4400/lep532 (C,G), and UT4400/lep754 (D,H) were added to monolayer cultured HeLa cells (1) and to RPMI2650 cells (2) at a cell to bacteria ratio of 1:100. They were allowed to react for 9 h and 6 h, respectively, and *E.coli* entry into the cells was observed under an electron microscope. Only UT4400/lep316 (B,F) was observed to invade HeLa cells and RPMI2650 cells. Although UT4400 (A,E), UT4400/lep532 (C,G) and UT4400/lep754 (D,H) were observed to be present around the cells, no invasion into the cells was observed with them. Magnification ×5000

### InvX, InvY, InvZ mediated invasion of *E.coli* into the nasal epithelial cells (gentamicin protection assay)

Next, using a gentamicin protection assay, the number of bacteria which entered into RPMI2650 cells was determined in colony forming units (CFU).

To determine uptake of the host *E.coli* cells using a gentamicin protection assay, we assessed the invasive ability of InvX, InvY, InvZ expressing *E.coli* (pMK100) cells showed invasion levels at the 3 h time point.

In RPMI2650 cells, invasive activity of InvX-expressing *E.coli* was significantly higher than that of InvY, InvZ, and negative control (Fig 5). The result was similar to the observations by electron microcopy. Invasion activity into nasal mucosa epithelial cells was successfully imparted to a pathogenic *E.coli* by externally expressing the InvX region of *M.leprae* on the *E.coli*. The InvX mediates the nasal epithelial cells invasion by non-pathogenic *E.coli*. The InvX region within mce1A protein is then sufficient for the invasion of *E.coli* into the cells.

**Fig 5.**
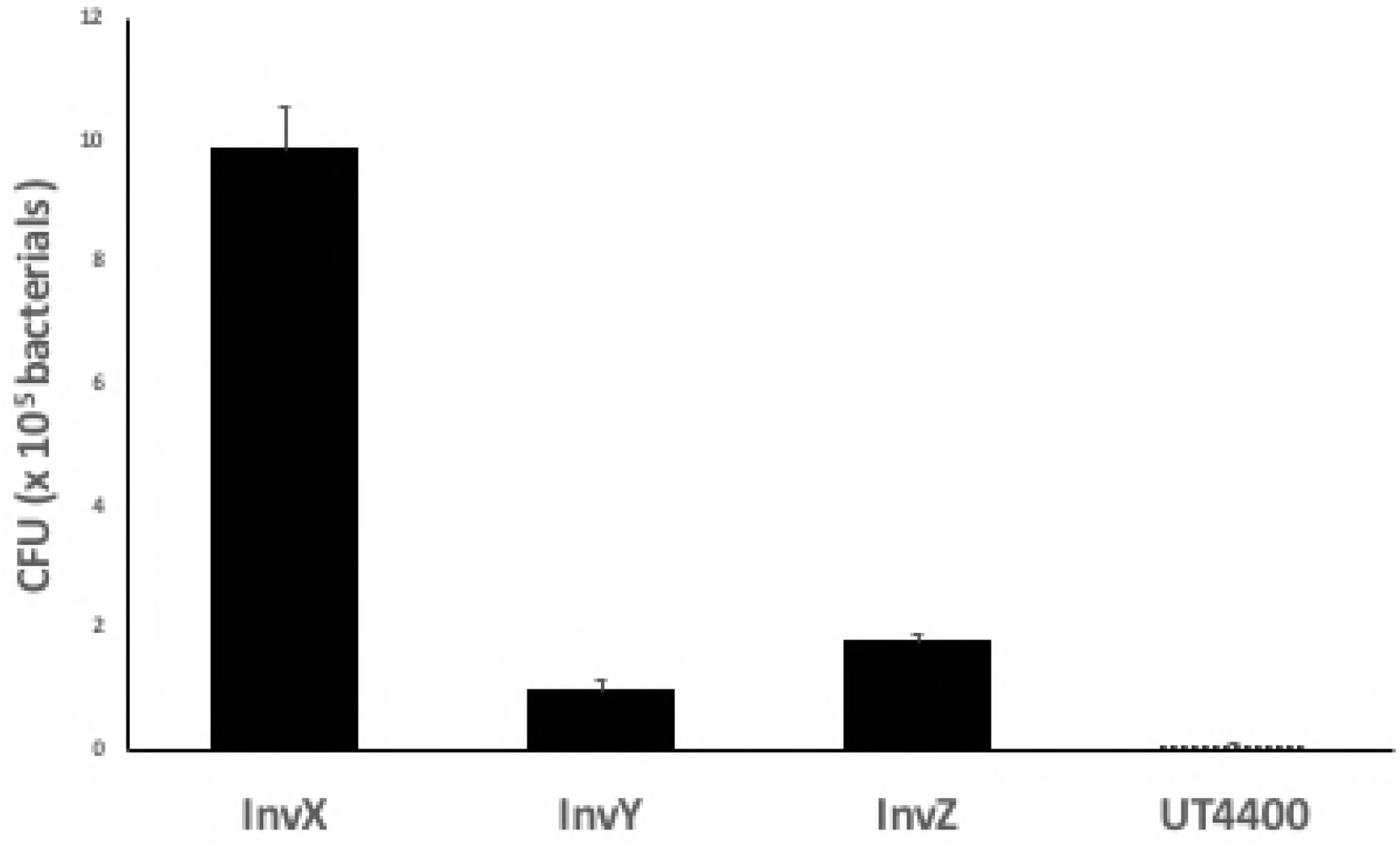
InvX, InvY, InvZ mediated invasion by *E.coli* into the nasal epithelial cells (gentamicin protection assay). Each of AIDA externally expressing *E.coli* was added to manolayer cultured RPMI2650 cells at a cells to bacteria ration of 1:10 and the mixture was allowed to react for 3 h. The bacteria outside the cell were subjected to disinfection by GM for 2 h. The bacteria inside the cell were counted with the colony count method, which was presented as CFU. Bars represent the mean ± S.D. of intracellular bacteria as a CFU in a representative experiment performed in triplicate. Asterisks indicate significant difference compared with an IgG control (Scheffe’s multiple comparison test).

### Indirect immunofluorescence staining of InvX expressing *E.coli* by antibodies corresponding to each region of mce1A

Indirect immunofluorescence was used to determine which regions of mce1A are sufficient to confer invasive ability to *E.coli*.

A 72-amino acid fragment of the InvX region was divided into four regions, InvXa (24 aa), InvXb (22 aa), InvXc (11 aa), and InvXd (15 aa). These peptides were subsequently synthetized as immune antigens for anti-InvX antibodies (anti-InvXa Ab, anti-InvXb Ab, anti-InvXc Ab, and anti-InvXd Ab) were studied.

In order to examine whether the antibodies recognize each of the regions, fluorescence immunostaining was conducted on the antibodies. The result was the following. Fluorescence microscopy revealed bacterial surface binding of the InvX antibodies by their binding of labelled secondary antibodies of fluorescence goat anti-rabbit IgG InvXa, InvXb, InvXc, and InvXd (Fig 6).

**Fig 6.**
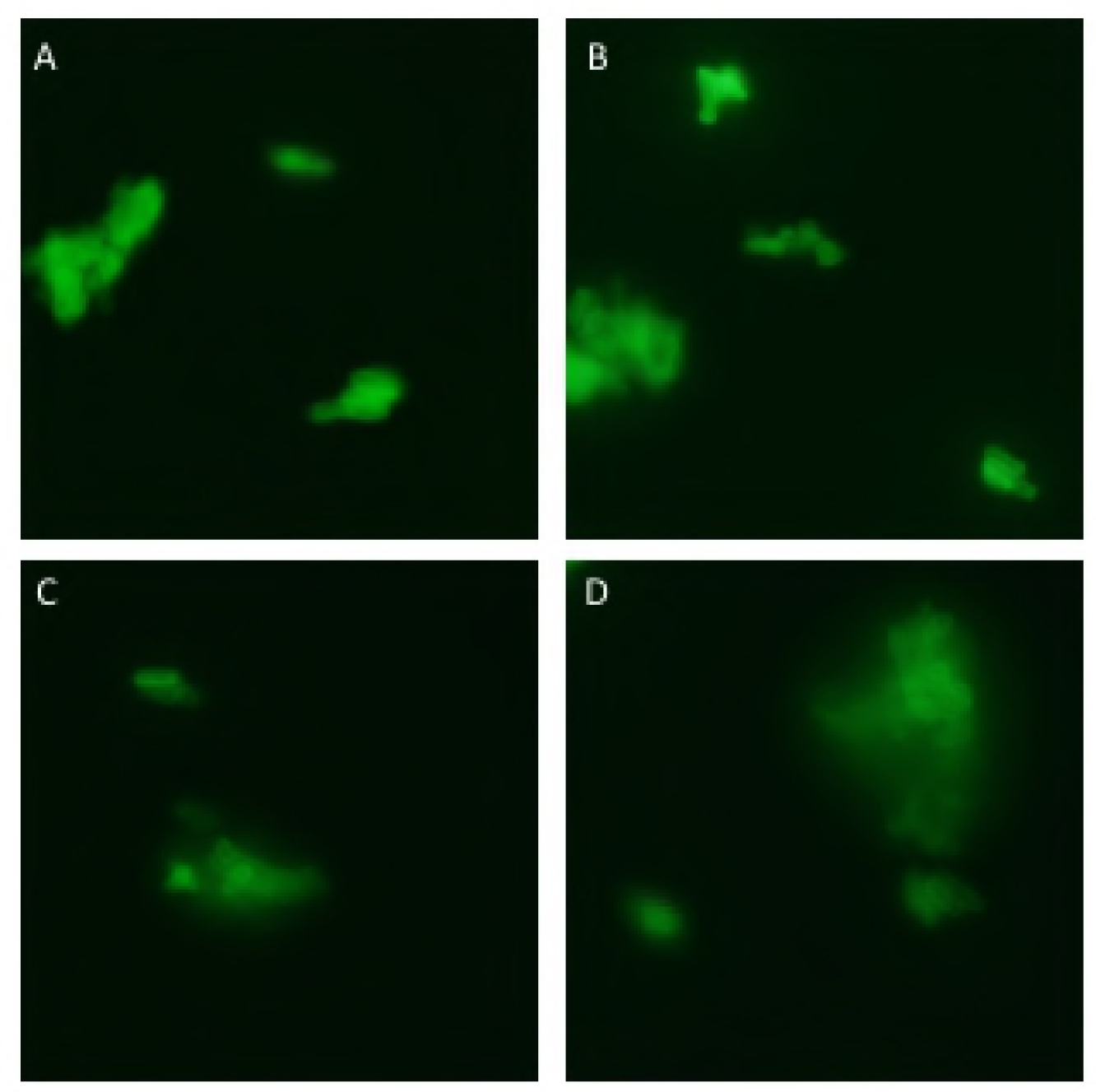
Indirect immunofluorescence staining of InvX expressing *E.coli* by antibodies corresponding to each region of InvX. Visualization of the bacilli reveals bacterial surface binding of the InvX antibody bound to fluorescent goat anti-rabbit IgG. (A) InvXa, (B) InvXb, (C) InvXc, (D) InvXd.

### Inhibitory effects of anti-InvX antibodies raised against each set of synthetic peptide corresponding to an InvX devided region on the nasal epithelial cells invasion of InvX expressing *E.coli*

In order to elucidate the role mce1A protein in association with the nasal epithelial cells invasion of *M.leprae*, we analyzed inhibitory effects of the resultant antibodies on the cell uptake of InvX-expressing *E.coli* by the inhibition assay. As shown in CFU analysis, the InvX-expressing *E. coli* pretreated with anti-InvXa Ab, anti-InvXb Ab, and anti-InvXd Ab had significantly lower entry than the IgG control, but there was no significant difference in pretreatment with anti-InvXc Ab and IgG control (Fig 7). These findings suggest that the invasion activity was most suppressed when using antibodies to cover the polypeptide chain encoded by 316 – 387 bp and expressed on the surface of *E.coli*.

**Fig 7.**
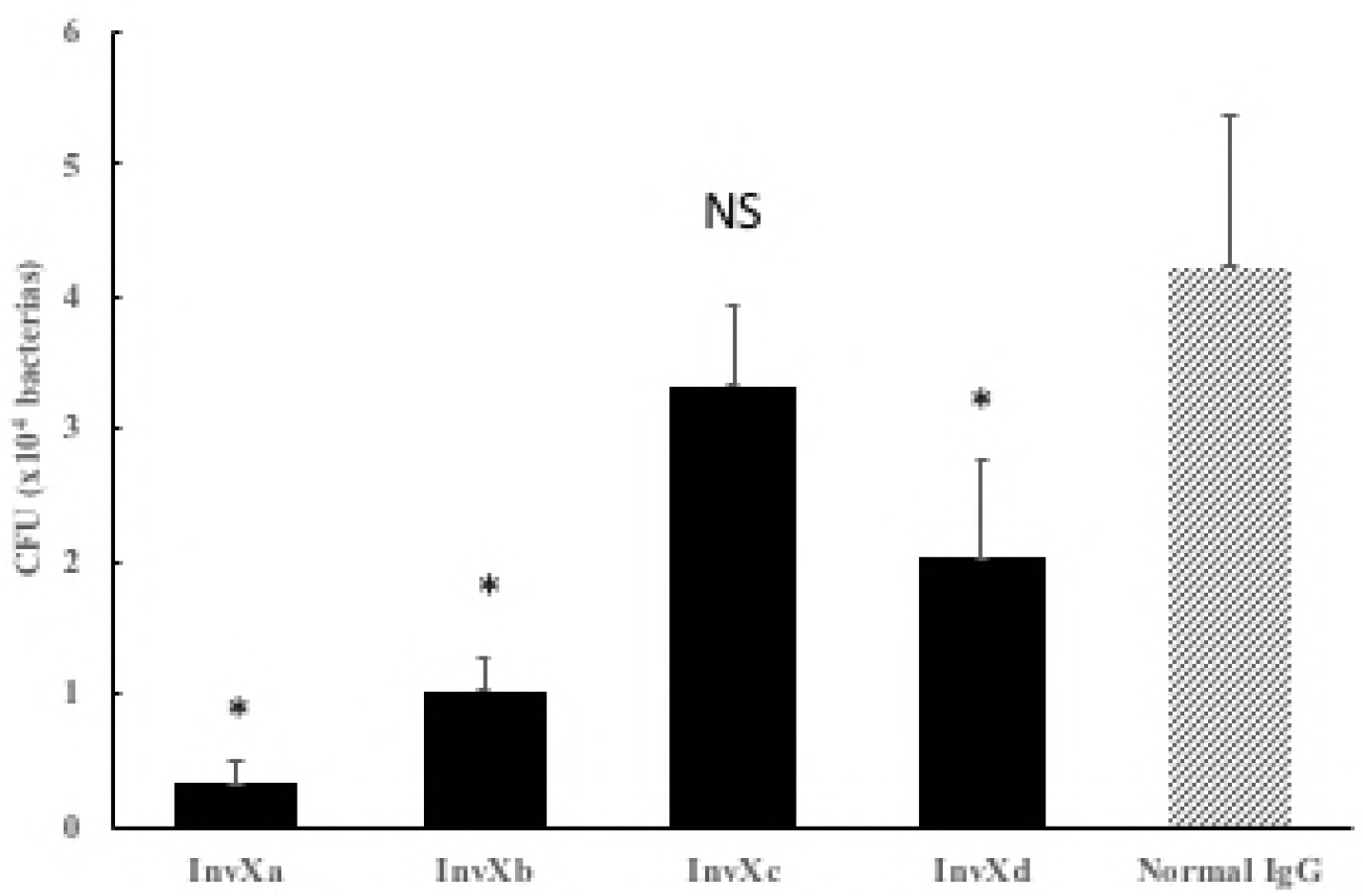
Inhibitory effects on anti-InvX antibodies raised against each set of synthetic peptide corresponding to an InvX divided region on the nasal epithelial cells invasion by InvX epressing *E.coli*. pMK100 *E.coli* strain were pretreated overnight with resultant antibodies and normal IgG as a control. Each antibody preparation was added to monolayer RPMI2650 cells and incubated for 3 h. After washing, GM was added to eliminate bacteria outside the cells. Bars represent mean ± standard deviation CFU of pMK100 *E.coli* treated with antibodies. The invasive activity was suppressed by antibodies against InvXa, InvXb, and InvXd. Asterisks indicate *p*<0.05, NS: Not significant (Dunnet test). Bars represent the mean ± SD of intracellular bacteria in CFU in a representative experiment performed in triplicate. Asterisks indicate a significant difference compared with an IgG control.

## Discussion

A number of studies have been conducted on the infection mode of *M.leprae*. In 1955, Khanolkar et al. reported that *M.leprae* infection of *M.leprae* occurs by normal skin contact[3]. However, in 1963 Weddell et al. revealed that the infection does not occur unless the bacteria is inoculated under the skin[24]. Rees et al. induced immune suppressed mice to inhale an aerosols containing *M.leprae* which successfully infected the mice via upper airway[4]. Following this, Chehl et al. revealed that transnasal infections of *M.leprae* of nude mice was possible[5]. From these studies, it became clear that the infection from aerosol containing *M.leprae* and through the nasal membrane can be established. However, as of today, only limited studies have been conducted regarding molecular mechanisms involved in the invasion.

The mce region is present in tuberculosis complex such as *M.tuberculosis* and *Mycobacterium bovis*, as well as in atypical mycobacteria such as *Mycobacterium avium* and *Mycobacterium intracellulare*[25]. Chitale et al. revealed that this mce1A protein involved in the invasion into epithelial cells is expressed only in tuberculosis complex[16]. We had found that *M.leprae* has a region highly homologous with to the mce1A region of *M.tuberculosis*, and so far have prepared a recombinant protein (mce1A protein) encoded by mce1A region (ML2589; 3092446 to 3093771, 1326 bp) of *M.leprae* to investigate invasion activities to epithelial cells[15,19]. In the present study, we have confirmed that the mce1A protein was actually expressed, as a native protein, on the surface of *M.leprae*, and prepared a recombinant protein by truncating the N terminus and C terminus of mce1A region of *M.leprae* to investigate the invasion activity into the epithelial cells. As a result, it was found that invasion activity is maintained even if 105 aa (315 bp) is truncated from N terminus and 308 aa (922 bp) is truncated from C terminus. Next, 316 bp to 921 bp region was divided into 3 parts, and each part was incorporated into an AIDA vector, where each region was externally expressed as a polypeptide chain to investigate whether the ability to invade can be imparted to non-pathogenic *E.*coli. These *E.coli* which externally express the protein by AIDA method were examined for the invasion activity using RPMI2650 cells, where the results indicate that active sequence of *M.leprae* involved in the invasion into nasal mucosa epithelial cells is present in the 316 – 531 bp of mce1A region.

The most important region of mce1A protein involved in the invasion of *M.tuberculosis* into human epithelial cells is called the InvIII cell and this is located between amino acids of position 130 to position 152[26]. The InvIII region of *M.tuberculosis* corresponds to InvXb of *M.leprae*. The sequence of the regions are identical between amino acid of position 10 to position 22 counted from N terminus, except that amino acids at positions 1 to 3, 5, 8, 9, 13 are different between *M.leprae* and *M.tuberculosis.* Suppression test results also indicated that the most important region of mce1A protein of *M.leprae* involved in the invasion into human epithelial cells is different from that of *M.tuberculosis.*

While *M.tuberculosis* has 3,959 protein-encoding genes and only 6 psedogenes, *M.leprae* has only 1,604 protein-encoding genes but has 1,116 pseudogenes[27], indicating that in *M.leprae*, far more proteins are inactivated as compared to *M.tuberculosis*. As in *M.tuberculosis*, the mce1A protein is expressed on the surface of bacteria as a native protein, and therefore the protein is considered as one of the most important proteins involved in the invasion of *M.leprae* into nasal mucosa epithelial cells.

## Acknowledgement

We thank Ms. Noriko Nemoto for excellent technical advice, and the staff of the Kitasato University Electron Microscope Center for technical assistance.

## References

1. WHO. Global leprosy update, 2016: accelerating reduction of disease burden. Wkly Epidemiol Rec. 2017 Sep 1; 92: 501–20.

2. Britton WJ, Lockwood DN. Leprosy. Lancet. 2004;363(9416):1209–19. Epub 2004/04/15. doi: 10.1016/S0140-6736(04)15952-7. PubMed PMID: 15081655.

3. Noordeen, SK. The epidemiology of leprosy. 2^nd^ ed. Edinburgh: Churchill-livingstone. 1994. pp. 29–48.

4. Rees RJ, McDougall AC. Airborne infection with *Mycobacterium leprae* in mice. J Med Microbiol. 1977;10(1):63–8. Epub 1977/02/01. doi: 10.1099/00222615-10-1-63. PubMed PMID: 320339.

5. Chehl S, Job CK, Hastings RC. Transmission of leprosy in nude mice. Am J Trop Med Hyg. 1985;34(6):1161–6. Epub 1985/11/01. PubMed PMID: 3914846.

6. McDougall AC, Rees RJ, Weddell AG, Kanan MW. The histopathology of lepromatous leprosy in the nose. J Pathol. 1975;115(4):215–26. Epub 1975/04/01. doi: 10.1002/path.1711150406. PubMed PMID: 1099180.

7. Job CK, Jayakumar J, Kearney M, Gillis TP. Transmission of leprosy: a study of skin and nasal secretions of household contacts of leprosy patients using PCR. Am J Trop Med Hyg. 2008;78(3):518–21. Epub 2008/03/14. PubMed PMID: 18337353.

8. Suneetha S, Arunthathi S, Job A, Date A, Kurian N, Chacko CJ. Histological studies in primary neuritic leprosy: changes in the nasal mucosa. Lepr Rev. 1998;69(4):358–66. Epub 1999/02/03. PubMed PMID: 9927808.

9. Patrocinio LG, Goulart IM, Goulart LR, Patrocinio JA, Ferreira FR, Fleury RN. Detection of *Mycobacterium leprae* in nasal mucosa biopsies by the polymerase chain reaction. FEMS Immunol Med Microbiol. 2005;44(3):311–6. Epub 2005/05/24. doi: 10.1016/j.femsim.2005.01.002. PubMed PMID: 15907454.

10. Klatser PR, van Beers S, Madjid B, Day R, de Wit MY. Detection of *Mycobacterium leprae* nasal carriers in populations for which leprosy is endemic. J Clin Microbiol. 1993;31(11):2947–51. Epub 1993/11/01. PubMed PMID: 8263180; PubMed Central PMCID: PMCPMC266167.

11. Rambukkana A, Salzer JL, Yurchenco PD, Tuomanen EI. Neural targeting of *Mycobacterium leprae* mediated by the G domain of the laminin-α2 chain. Cell. 1997;88(6):811–21. Epub 1997/03/21. PubMed PMID: 9118224.

12. Rambukkana A, Yamada H, Zanazzi G, Mathus T, Salzer JL, Yurchenco PD, et al. Role of α-dystroglycan as a Schwann cell receptor for *Mycobacterium leprae*. Science. 1998;282(5396):2076–9. Epub 1998/12/16. PubMed PMID: 9851927.

13. Shimoji Y, Ng V, Matsumura K, Fischetti VA, Rambukkana A. A 21-kDa surface protein of *Mycobacterium leprae* binds peripheral nerve laminin-2 and mediates Schwann cell invasion. Proc Natl Acad Sci U S A. 1999;96(17):9857–62. Epub 1999/08/18. PubMed PMID: 10449784; PubMed Central PMCID: PMCPMC22300.

14. Ng V, Zanazzi G, Timpl R, Talts JF, Salzer JL, Brennan PJ, et al. Role of the cell wall phenolic glycolipid-1 in the peripheral nerve predilection of *Mycobacterium leprae*. Cell. 2000;103(3):511–24. Epub 2000/11/18. PubMed PMID: 11081637.

15. Arruda S, Bomfim G, Knights R, Huima-Byron T, Riley LW. Cloning of an *M. tuberculosis* DNA fragment associated with entry and survival inside cells. Science. 1993;261(5127):1454–7. Epub 1993/09/10. PubMed PMID: 8367727.

16. Chitale S, Ehrt S, Kawamura I, Fujimura T, Shimono N, Anand N, et al. Recombinant *Mycobacterium tuberculosis* protein associated with mammalian cell entry. Cell Microbiol. 2001;3(4):247–54. Epub 2001/04/12. PubMed PMID: 11298648.

17. Casali N, Konieczny M, Schmidt MA, Riley LW. Invasion activity of a Mycobacterium tuberculosis peptide presented by the *Escherichia coli* AIDA autotransporter. Infect Immun. 2002;70(12):6846–52. Epub 2002/11/20. PubMed PMID: 12438361; PubMed Central PMCID: PMCPMC133103.

18. Fujimura T. Lu S, Ehrt S, Shimono N, Riley LW. Examination of anti-mcep monoclonal antibodies in their inhibitory effect on mammalian cell association. In the US-Japan cooperative medical science program. 34th Tuberculosis-Leprosy research conference. San Francisco: California. 1999 Jun 27-30. pp. 271.

19. Sato N, Fujimura T, Masuzawa M, Yogi Y, Matsuoka M, Kanoh M, et al. Recombinant *Mycobacterium leprae* protein associated with entry into mammalian cells of respiratory and skin components. J Dermatol Sci. 2007;46(2):101–10. Epub 2007/02/24. doi: 10.1016/j.jdermsci.2007.01.006. PubMed PMID: 17317107.

20. Maeda S, Matsuoka M, Nakata N, Kai M, Maeda Y, Hashimoto K, et al. Multidrug resistant *Mycobacterium leprae* from patients with leprosy. Antimicrob Agents Chemother. 2001;45(12):3635–9. Epub 2001/11/16. doi: 10.1128/AAC.45.12.3635-3639.2001. PubMed PMID: 11709358; PubMed Central PMCID: PMCPMC90887.

21. Matsuoka M. The history of *Mycobacterium leprae* Thai-53 strain. Lepr Rev. 2010;81(2):137. Epub 2010/09/10. PubMed PMID: 20825118.

22. Benz I, Schmidt MA. Cloning and expression of an adhesin (AIDA-I) involved in diffuse adherence of enteropathogenic *Escherichia coli*. Infect Immun. 1989;57(5):1506–11. Epub 1989/05/01. PubMed PMID: 2565291; PubMed Central PMCID: PMCPMC313306.

23. Elsinghorst EA. Measurement of invasion by gentamicin resistance. Methods Enzymol. 1994;236:405–20. Epub 1994/01/01. PubMed PMID: 7968625.

24. Weddell G, Palmer E, Rees RJW, Jamison DC. Experimental observations related to the histopathology of leprosy. In the pathogenesis of Leprosy. Ciba Foundations Study Group No 15 (eds). London: Churchill. 1963. pp. 3–15.

25. Parker SL, Tsai YL, Palmer CJ. Comparison of PCR-generated fragments of the mce gene from *Mycobacterium tuberculosis*, *M. avium*, *M. intracellulare*, and *M. scrofulaceum*. Clin Diagn Lab Immunol. 1995;2(6):770–5. Epub 1995/11/01. PubMed PMID: 8574846; PubMed Central PMCID: PMCPMC170237.

26. Kohwiwattanagun J, Kawamura I, Fujimura T, Mitsuyama M. Mycobacterial mammalian cell entry protein 1A (Mce1A)-mediated adherence enhances the chemokine production by A549 alveolar epithelial cells. Microbiol Immunol. 2007;51(2):253–61. Epub 2007/02/21. PubMed PMID: 17310094.

27. Cole ST, Eiglmeier K, Parkhill J, James KD, Thomson NR, Wheeler PR, et al. Massive gene decay in the leprosy bacillus. Nature. 2001;409(6823):1007–11. Epub 2001/03/10. doi: 10.1038/35059006. PubMed PMID: 11234002.

